# Detection and phenotypic characterization of carbapenem non susceptible gram-negative bacilli isolated from clinical specimens

**DOI:** 10.1101/2021.08.17.456671

**Authors:** Abera Abdeta, Adane Bitew, Surafel Fentaw, Estifanos Tsige, Dawit Assefa, Tadesse Lejisa, Yordanos Kefyalew, Eyasu Tigabu

**Affiliations:** National Clinical Bacteriology and Mycology Reference Laboratory, Ethiopian public health institute, Ethiopia; Department of Medical Laboratory Sciences, College of Health Sciences, Addis Ababa University, Addis Ababa, Ethiopia; National Clinical Chemistry Reference Laboratory, Ethiopian public health institute, Ethiopia; Department of Applied Biology, School of Applied Natural Science, Adama Science and Technology University; Global One Health initiative, The Ohio State University, East African Regional Office

## Abstract

**Background:** Multi-drug resistant, extremely drug-resistant, pan-drug resistant, carbapenem-resistant, and carbapenemase-producing gram-negative bacteria are becoming more common in health care settings and are posing a growing threat to public health.

**Objective:** The study was aimed to detect and phenotypically characterize carbapenem non susceptible gram-negative bacilli at Ethiopian Public Health Institute.

**Materials and methods:** Prospective cross-sectional study was conducted from June 30, 2019, to May 30, 2020, at the national reference laboratory of the Ethiopian Public Health Institute. Clinical samples were collected, inoculated, and incubated in accordance to standard protocol for each sample. Antimicrobial susceptibility testing was done using Kirby Bauer disk diffusion. Identification was done using the traditional biochemical method. Multidrug-resistant and extensively drug-resistant were classified using a standardized definition established by European Centers for Disease prevention and control and the United States Centers for Disease prevention and control experts. Carbapenemase production was confirmed by modified carbapenem inactivation and a simplified carbapenem inactivation method. Meropenem with EDTA was used to differentiate serine carbapenemase and Metallo β-lactamase.

**Results:** A total of 1337 clinical specimens were analyzed, of which 429-gram negative bacilli isolates were recovered. Out of 429 isolates 319, 74, and 36 were Enterobacterales, *Acinetobacter* species, and *P. aeruginosa* respectively. In our study, the prevalence of Multidrug-resistant, extensively drug-resistant, Carbapenemase-producing, and carbapenem non-susceptible Gram-negative bacilli were, 45.2%, 7.7%, 5.4%, and 15.4% respectively. Out of 66 isolates screened for Carbapenemase, 34.8% (23/66) were Carbapenemase enzyme producers. Ten out of twenty-three Carbapenemase-positive organisms were Metallo-beta-lactamase producers. Thirteen out of twenty-three isolates were serine carbapenemase producers. Three out of 13 serine Carbapenemase positive organisms were *Klebsiella pneumoniae* Carbapenemase.

**Conclusion:** The finding from this study revealed a high prevalence of Multidrug-resistant, extremely drug-resistant, carbapenemase-producing gram-negative bacteria, particularly among Intensive care unit patients at the health facility level, this necessitates a robust laboratory-based antimicrobial resistance monitoring and infection prevention and control program.

## Introduction

The modern discovery of the antimicrobial agent is considered one of the most fundamental milestones accomplished in the history of medicine and the discovery saved millions of lives [1]. Antimicrobials were first used to treat infections in the 1940s [2]. Shortly after the discovery of, Antimicrobials, the fast spread of antimicrobial resistance poses a serious threat to global public health [2].

The extensive use of Antimicrobials for treating humans and animal infections along with globalization and international travel leads to the fast spread of resistant strains globally [3]. Particularly, increasing incidence of healthcare-associated infections due to multi-drug resistant (MDR), extremely drug-resistant (XDR), carbapenem-resistant, and Carbapenemase-producing Gram-negative bacilli (GNB) has been widely reported [4-7].

International subject matter experts came together through a joint initiative by the European Centre for Disease Prevention and Control (ECDC) and the United States Centers for Disease Control and Prevention (CDC), to create a standardized international definition with which to describe acquired resistance profiles [8].

They defined MDR as acquired non-susceptibility to at least one agent in three or more antimicrobial categories, XDR as non-susceptibility to at least one agent in all but two or fewer antimicrobial categories (i.e., bacterial isolates remain susceptible to only one or two categories), and PDR as non-susceptibility to all agents in all antimicrobial categories. To apply these definitions, bacterial isolates should be tested against all or nearly all of the antimicrobial agents within the antimicrobial categories and selective reporting and suppression of results should be avoided [8].

The common mechanism of developing resistance to carbapenem antibiotics is through carbapenemase enzyme production [9]. Carbapenemase is the most versatile family of β lactamase and recognizes almost all hydrolyzable β lactams, and most are resilient against inhibition by all available β lactamase inhibitors [9]. *Klebsiella pneumonia* Carbapenemase (KPC) hydrolyze penicillin, all cephalosporins, monobactams, carbapenems, and β lactamase inhibitors. Metallo-β-lactamases usually exhibit resistance to penicillin, cephalosporins, carbapenems, and the clinically available β lactamase inhibitors but are inhibited by monobactams [10].

There is a considerable knowledge gap regarding risk factors associated with the occurrence of MDR bacteremia [11]. Identifying risk factors for acquiring gram-negative bacteremia could potentially help patient care and management [11].

Despite the growing global burden of multidrug resistance, extremely drug-resistant, and carbapenemase-producing gram-negative bacilli, data on multidrug resistance, extremely drug resistance, and carbapenemase-producing organisms that includes both fermentative and non-fermentative gram-negative bacilli in Ethiopia is scarce. Hence, the objective of this study was to determine the prevalence of MDR, XDR, carbapenem non-susceptible, and carbapenemase-producing gram-negative bacilli isolated from various clinical specimens, as well as to phenotypically characterize carbapenem non-susceptible isolates, at the Ethiopian Public Health Institute,

## Materials and methods

### Study design, site, and period

A prospective cross-sectional study was conducted from June 30, 2019, to May 30, 2020, at the National Clinical Bacteriology and Mycology Reference Laboratory on clinical samples collected there and referred from different health care settings in Addis Ababa.

### Sample collection and processing

Microbiological specimens were collected from the body fluids, ear swabs, sputum, urine, pus, cerebrospinal fluid, blood, and tracheal aspirates and processed following standard procedures. In case sample transportation delay is inevitable appropriate transport media were used. A total of 1337 clinical specimens were collected during the study period. Specimens collected from each patient were inoculated onto culture media and incubated at appropriate temperatures and periods according to standard protocols related to each sample. Identification was done using the conventional biochemical method. Gram staining, colony characterization, and biochemical tests were done as part of identification. AST was done by Kirby Baur disk diffusion method as per CLSI M100 2020. All frequently isolated Enterobacterales, *Acinetobacter* species, and *P. aeruginosa* recovered from the various clinical specimen during the study period were included. All the necessary variables such as socio-demographic (age and sex), type of specimen, referring health facilities, patient location, previous antibiotic exposure from the request form were entered onto pre-configured WHOnet software version 2019.

### Bacterial identification and antimicrobial susceptibility testing

#### Bacterial identification

The samples were inoculated onto appropriate culture media, incubated at appropriate temperature and time following standard procedure. The growth was inspected to identify the bacteria. Presumptive identification of bacteria was done based on Gram reaction and colony characteristics. Confirmatory tests were done by enzymatic and biochemical properties of the organisms for the final identification of the isolates. Gram-negative rods were identified by performing a series of biochemical tests such as carbohydrate fermentation on triple sugar iron agar, Oxidase strips, Simon’s citrate agar, and lysine iron agar. Indole production and motility were checked on the sulfide-indole-motility (SIM) medium. Urease production was inspected using a urea agar base supplemented with 40% urea solution.

#### Antimicrobial susceptibility testing

Kirby Bauer disk diffusion method was used on Muller Hinton agar to determine antimicrobial susceptibility patterns of the isolates and CLSI M100 2020 was used to interpret the results [12].

The Antimicrobial discs used for Kirby Bauer disk diffusion method were in the following concentrations: Ampicillin(10µl), amoxicillin/clavulanic acid(20/10µl), piperacillin/tazobactam(1 00/10µl), cefazolin(30µl), cefuroxime(30µl), ceftazidime(30µl), ceftriaxone(30µl), cefotaxime(3 0µl), cefepime(30µl), imipenem(10µl), meropenem(10µl), amikacin(30µl), gentamicin(10µl), tobramycin(10µl), nalidixic acid(30µl), ciprofloxacin(5µl), trimethoprim/sulfamethoxazole (1.25/23.75µl), nitrofurantoin(300µl), and tetracycline(30µl).

### Detection of Carbapenemase

Clinical and Laboratory Standards Institute CLSI (2010) introduced the modified Hodge test for Carbapenemase detection, but this method can only be used for the accurate detection of KPC-type Carbapenemase in Enterobacteriaceae [13]. CLSI (2012) recommended the Carba NP test method for the detection of Carbapenemase in gram-negative bacilli; however, the preparation of the reagents required for this test is complicated and the solutions cannot be stored for extended periods, limiting its clinical application [14].

In 2015 a new detection method, the carbapenem inactivation method (CIM), which is easy to operate and highly sensitive in the detection of Carbapenemase was designed [15]. In 2017, based on the CIM method, CLSI recommended the modified carbapenem inactivation method (mCIM), However; it is a relatively complex method and can only be used to detect Carbapenemase in Enterobacteriaceae and *P. aeruginosa* [16]. In 2018, based on the mCIM, a simplified carbapenem inactivation method (sCIM) was designed for simple and accurate detection of Carbapenemase in gram-negative bacilli [17].

### Modified Carbapenem Inactivation method

In the mCIM, 1 mL loop full of Enterobacteriaceae or 10 mL loop full of *P. aeruginosa* from blood agar plates was emulsified in 2 mL trypticase soy broth (TSB). A meropenem disk was then immersed in the suspension and incubated for a minimum of 4 h at 35°C. A 0.5 McFarland suspension of *E. coli* ATCC 25922 was prepared in saline using the direct colony suspension method. A Mueller Hinton agar (MHA) plate was inoculated with *E. coli* ATCC 25922 using the routine disk diffusion procedure. The meropenem disk was removed from the TSB and placed on an MHA plate previously inoculated with the *E. coli* ATCC 25922 indicator strains. Plates were incubated at 35°C in ambient air for 18-24 h. An inhibition zone diameter of 6-15 mm or colonies within a 16–18 mm zone was considered to be a positive result, and a zone of inhibition of ≥19 mm was considered to be a negative result [12].

### Simplified Carbapenem Inactivation Method

The sCIM is based on the mCIM with an improvement of experimental procedures. Instead of incubating the antimicrobial disk in the organism culture media for 4 hours as in the mCIM, the organism to be tested was smeared directly onto an antimicrobial disk in the sCIM. To perform the sCIM, for *Acinetobacter* species, a 0.5 McFarland standard suspension (using direct colony suspension method) of *E. coli* ATCC 25922 was diluted 1:10 in saline and inoculated onto the MHA plate, following the routine disk diffusion procedure. Plates were allowed to dry for 3–10 min [17].

Then, 1–3 overnight colonies of the test organisms grown on blood agar were smeared onto one side of an imipenem disk (10µg); immediately afterward, the side of the disk having bacteria was placed on the MHA plate previously inoculated with *E. coli* ATCC 25922. An imipenem disk placed on an MHA plate was used as the control [17].

All plates were incubated at 35C for 16–18 h in ambient air. Bacterial strains that produced Carbapenemase can hydrolyze imipenem; hence the susceptible indicator strain grew unchecked. Zone of inhibition around the disk shows a diameter of 6–20 mm, or the satellite growth of colonies of *E. coli* ATCC 25922 around the disk with a zone diameter ≤22 mm, was considered carbapenemase positive; a zone of inhibition ≥26 mm was considered to be a negative result; a zone of inhibition of 23–25 mm was considered to be a carbapenemase indeterminate result [17].

### Differentiation of Metallo β-lactamase from serine Carbapenemases

The Modified carbapenem inactivation method positive Enterobacterales (formerly Enterobacteriaceae) and *P. aeruginosa* and Simplified Carbapenem Inactivation Method positive *Acinetobacter* species detected were further screened for Class B Metallo-carbapenemase (MBLs), which are characterized by inhibition by metal chelators, EDTA. A ≥ 5-mm increase in zone diameter for eCIM vs. zone diameter for mCIM was considered a Metallo carbapenemase producer. A≤4mm increase in zone diameter for the eCIM vs zone diameter of mCIM was considered Metallo-carbapenemase negative. Carbapenemase positive, Metallo carbapenemase negative gram-negative bacilli were considered serine carbapenemase producers [12]

Quality control recommendation: *K. pneumoniae* ATCC® BAA-1705™ *E. coli* ATCC® 25922™ was used to check the quality of meropenem with EDTA [12].

### Differentiation of *Klebsiella pneumoniae* carbapenemase (KPC) from other serine Carbapenemases

Serine carbapenemase producers were further screened for *Klebsiella pneumoniae* Carbapenemase (KPC). MIC Test Strip KPC strips consisting of Meropenem (MRP)/Meropenem+Phenylboronic acid (MBO) were used to detect *Klebsiella pneumoniae* Carbapenemase (KPC) producing gram-negative. [18].

Well, isolated colonies from an overnight blood agar plate were suspended into saline to achieve a 0.5 McFarland standard turbidity (1 McFarland if mucoid) to obtain a confluent lawn of growth after incubation. The strip was applied to the agar surface with the scale facing upwards and the code of the strip to the outside of the plate. The agar plates were incubated in an inverted position at 35 ± 2°C for 16-20 hours in the ambient atmosphere. The incubation time was Extend up to 48 hours in case of slow-growing Gram-negative non-fermenters [18]

Result Interpretation: The ratio of MRP/MBO of ≥8 or ≥3 log2 dilutions was interpreted as KPC producer. The phantom zone or deformation of the ellipse was interpreted as positive for KPC regardless of the MRP/MBO ratio [18].

Quality control recommendations: *E. coli* ATCC® 25922 and ATCC® BAA-1705 (intrinsic KPC production) were used to check the quality of KPC strips [18].

## Data Quality Assurance

The quality of culture media, antimicrobial disks, and gradient strip were checked as per CLSI, EUCAST guideline, Laboratory SOPs, and manufacturer’s instructions.

## Data analysis and interpretation

The data were entered, cleaned, and analyzed using SPSS version 23 and WHOnet 2019 version. Tables and figures were used to present the results. Chi-square and Univariate analysis were used to determine the association between multidrug-resistant gram-negative bacilli and different risk factors. P-values less than 0.05 were considered statistically significant.

## Ethical considerations

The study was conducted after ethical clearance was obtained from the department research and ethical review committee of the Department of Medical Laboratory Sciences, College of Health Sciences, Addis Ababa University. Official permission from Ethiopian Public Health Institute was obtained. All results were kept confidential; the patient’s name and other personal identifiers were encrypted, rather the sample identification number automatically generated by Polytech was used.

## Results

During the study period, 1337 samples were analyzed, of which 429-gram negative isolates were recovered. Of which, 293, 74, and 36 were Enterobacterales, *Acinetobacter* species, and *P. aeruginosa* were respectively. The number of samples based on specimen types were as follows: Blood (364), Cerebrospinal fluid (46), Ear swabs (28), other body Fluids (30), Pus (366), Sputum (10), Stool (18), Tracheal aspirate (6), and Urine (469). Two hundred thirty-three and one hundred ninety-six-gram negative isolates were recovered from specimens collected from male and female patients respectively.

Most of the isolates were isolated from specimens referred from Aabet hospital (187), Ras Desta Hospital (94), and Saint peter Hospital (44). The distribution of gram-negative bacilli among health facilities is summarized in Figure 1.

**Figure 1:**
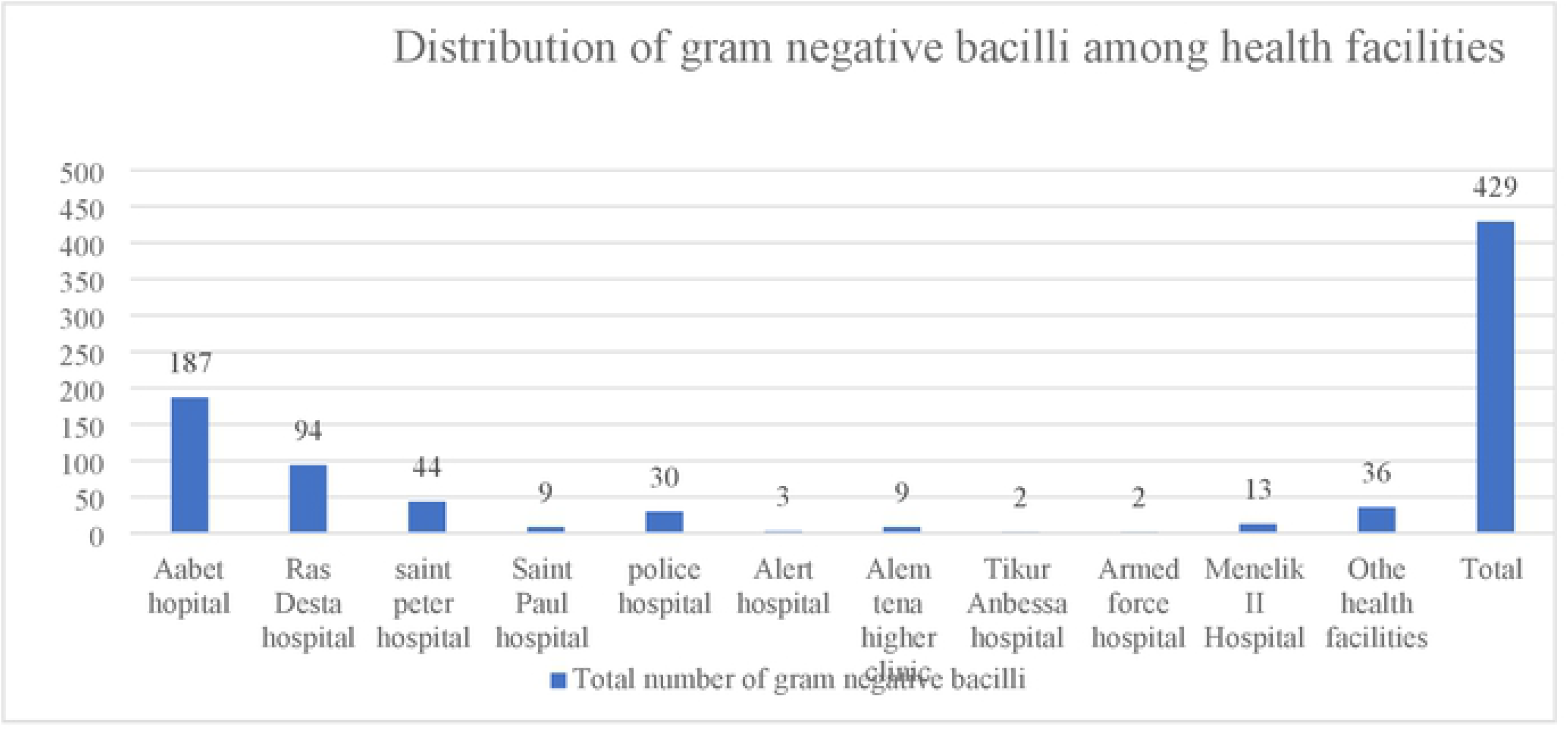
Distribution of gram-negative bacilli among health facilities.

The highest number of MDR, XDR, and carbapenemase-producing isolates were recovered from specimens referred from Aabet hospital, Ras Desta hospital, and Saint Peter hospital respectively. One hundred eighty-seven (187) gram-negative bacilli were recovered from Aabet hospital specimens, of which 130(69.5%), 15(8%), and 11(5.9%) were MDR, XDR, and Carbapenemase producers respectively. Ninety-four (94) gram-negative bacilli were recovered from Ras Desta hospital specimens, of which 40(42.5%),10(10.6%), and 7(7.4%) were MDR, XDR, and Carbapenemase producers, respectively. The distribution of MDR, XDR, and carbapenemase-producing isolates among health facilities was summarized in Figure 2.

**Figure 2:**
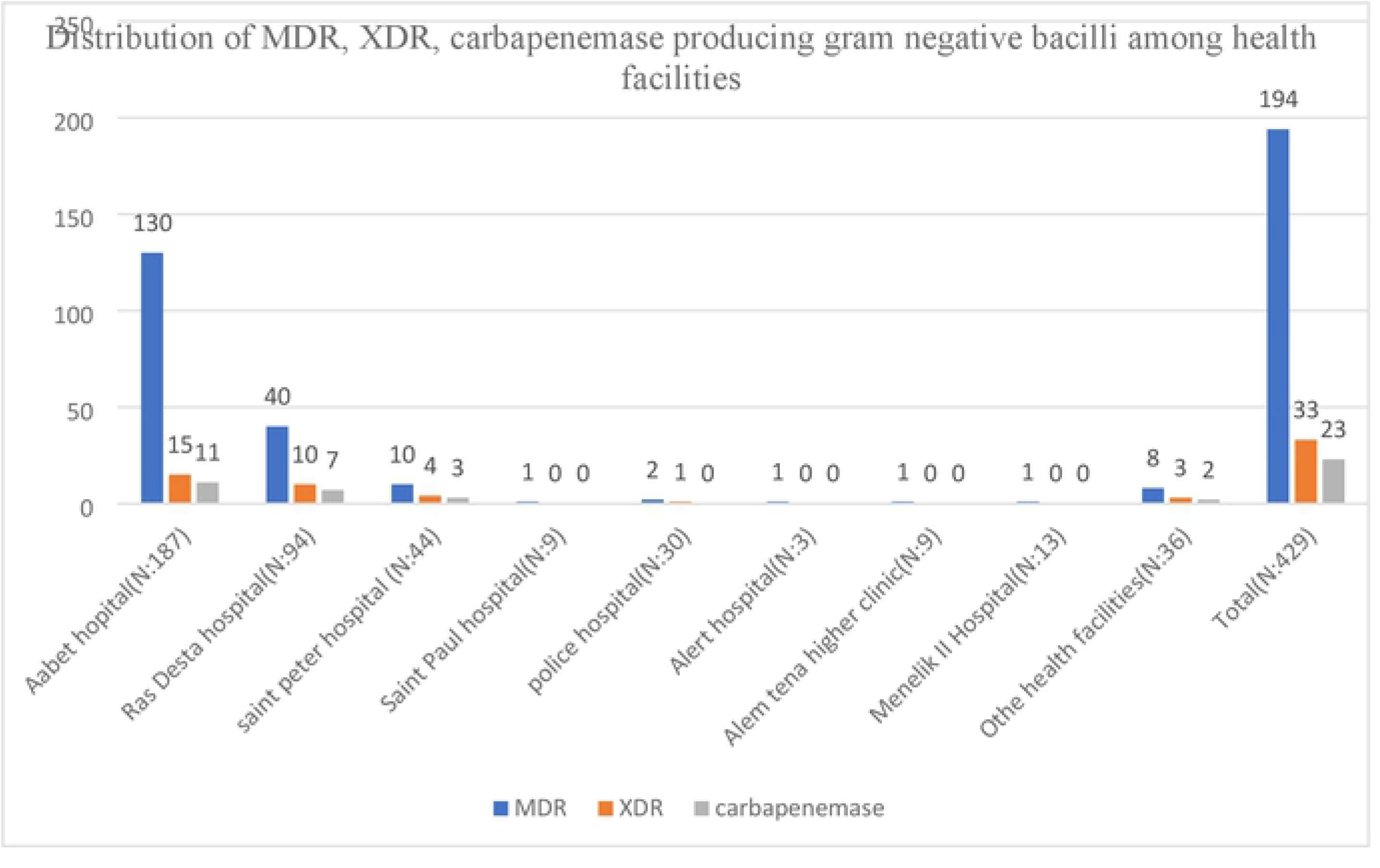
Distribution of MDR, XDR, carbapenemase-producing gram-negative bacilli among health facilities.

Out of four hundred twenty-nine (429) gram-negative bacilli, clinical isolates analyzed, (N=33,7.7%) were extremely drug-resistant. Out of 33 extremely drug-resistant gram-negative isolates; *Acinetobacter* species were the predominant isolates 72.73%[N=24]. The remaining nine XDR isolates were as follows; *K. Pneumonia* 9.09%[N=3], *P. mirabilis* 12.12%[N=4], *E. cloacae* 3.03%[N=1], and *P. aeruginosa* 3.03%[N=1] Table 1. Out of 33 XDR isolates [N=21,63.64%] were isolated from intensive care units. The highest number of XDR gram-negative bacilli were isolated from urine samples [N=16, 48.48%]. The distribution of XDR isolates, among specimen types, health facility wards were summarized in Table 1.

**Table 1.** Distribution of Extremely drug-resistant gram-negative bacilli.

| Distribution of XDR GNB |  |  | XDR GNB from different units |  |  | XDR GNB based on specimen sources |  |  |
| --- | --- | --- | --- | --- | --- | --- | --- | --- |
| Organism name | N | % | Ward | N | % | Specimen types | N | % |
| *Acinetobacter* species | 24 | 72.73 | Intensive care unit | 21 | 63.64 | Urine | 16 | 48.48 |
| *K. Pneumonia* | 3 | 9.09 | Burn unit | 2 | 6.06 | Blood | 7 | 21.21 |
| *P. mirabilis* | 4 | 12.12 | Orthopedics | 3 | 9.09 | Sputum | 1 | 3.03 |
| *E. cloacae* | 1 | 3.03 | Unknown | 6 | 18.18 | CSF | 1 | 3.03 |
| *P. aeruginosa* | 1 | 3.03 | Emergency | 1 | 3.03 | pus | 8 | 24.24 |
| *Total* | 33 | 100 | *Total* | 33 | 100 | *Total* | 33 | 100 |
| XDR: Extremely drug-resistant, GNB: Gram-negative bacilli: N: Number, %: percentage |  |  |  |  |  |  |  |  |

Out of 429 isolates, 194 were MDR GNB isolates. The most frequently isolated MDR organism was *K. Pneumoniae* [N=80, 73.4%] followed by *Acinetobacter* species [N=52,70.3%], and *E. coli* [N=36, 23.6%], Table 2. The remaining MDR GNB were summarized in Table 2

**Table 2.** Prevalence of multi-drug resistance and Extremely drug resistance, and carbapenemase-producing GNB against 8 antimicrobial classes.

| **Organisms** | **R<sub>0</sub>** | **R<sub>1</sub>** | **R<sub>2</sub>** | **R<sub>3</sub>** | **R<sub>4</sub>** | **R<sub>5</sub>** | **R<sub>6</sub>** | **R<sub>7</sub>** | **R<sub>8</sub>** | **N%**<br>**MDR** | **N%**<br>**XDR** | **N%**<br>**Carbapenemase** | **N%**<br>**MBL** | **N%**<br>**Serine** | **N%**<br>**KPC** |
| --- | --- | --- | --- | --- | --- | --- | --- | --- | --- | --- | --- | --- | --- | --- | --- |
| *E. coli*(N=152) | 3 | 52 | 61 | 20 | 14 | 2 | 0 | 0 | 0 | 36(23.7) | 0(0) | 2(1.3) | 1(0.7) | 1(0.7) | 1(0.7) |
| *K. pneumoniae*(N=109) | 0 | 0 | 36 | 51 | 10 | 9 | 0 | 1 | 2 | 80(73.4) | 3(2.8) | 9(8.3) | 5(4.6) | 4(3.7) | 1(0.9) |
| *K. oxytoca*(N=10) | 0 | 0 | 9 | 1 | 0 | 0 | 0 | 0 | 0 | 1(10) | 0(0) | 0(0) | 0(0) | 0(0) | 0(0) |
| *K. Ozaenae*(N=11) | 0 | 0 | 2 | 4 | 5 | 0 | 0 | 0 | 0 | 2(18.2) | 0(0) | 2(18.2) | 0(0) | 2(18.2) | 0(0) |
| *E. cloacae*(N=11) | 0 | 1 | 6 | 1 | 2 | 0 | 0 | 0 | 1 | 4(36.4) | 1(9.1) | 0(0) | 0(0) | 0(0) | 0(0) |
| *P. mirabilis*(N=17) | 0 | 1 | 10 | 1 | 1 | 0 | 0 | 2 | 2 | 6(35.3) | 4(23.5) | 1(5.9) | 1(5.9) | 0(0) | 0(0) |
| *P. vulgaris*(N=3) | 0 | 0 | 1 | 1 | 1 | 0 | 0 | 0 | 0 | 2(66.7) | 0(0) | 0(0) | 0(0) | 0(0) | 0(0) |
| *M.morganii* (N=6) | 0 | 0 | 5 | 0 | 1 | 0 | 0 | 0 | 0 | 1(16.7) | 0(0) | 0(0) | 0(0) | 0(0) | 0(0) |
| *Acinetobacter species*(N=74) | 0 | 0 | 22 | 0 | 3 | 5 | 20 | 10 | 14 | 52(70.3) | 24(32.4) | 5(6.8) | 2(2.7) | 3(4.1) | 0(0) |
| *P. aeruginosa*(N=36) | 0 | 3 | 23 | 5 | 4 | 0 | 0 | 0 | 1 | 10(27.8) | 1(2.8) | 4(11.1) | 1(2.8) | 3(8.4) | 1(2.8) |
| **Total(N=429)** | 3 | 57 | 175 | 84 | 41 | 16 | 20 | 13 | 20 | 194(45.2) | 33(7.7) | 23(5.4) | 10(2.3) | 13(3) | 3(0.7) |

Abbreviations: R<sub>0</sub>-R<sub>8</sub>: No resistant to antimicrobial class to Resistant to eight antimicrobial classes, MDR-Multi-drug resistant, XDR-Extensively drug-resistant, MBL-Metallo- β-lactamase, KPC-Klebsiella pneumoniae carbapenemase, N-Number, %-percentage.

Out of 429 isolates, 15.4% (66/429) isolates were non-susceptible to either meropenem or Imipenem. Carbapenem non-susceptible isolates were considered a candidate for carbapenemase screening. Out of 66 isolates screened for Carbapenemase, 34.8% (23/66) were Carbapenemase enzyme producers Table 2. Ten out of twenty-three Carbapenemase-positive organisms were Metallo-beta-lactamase (MBL) producers. Thirteen out of twenty-three isolates were serine carbapenemase producers. Three out of thirteen serine Carbapenemase-positive organisms were *Klebsiella pneumoniae* Carbapenemase (KPC). Out of 10 Metallo-beta-lactamase positive isolates, 5(50%) were *K. Pneumoniae*, and the remaining were summarized in Table 2. Three KPC isolates were, *E. coli, K. Pneumoniae*, and *P. aeruginosa* Table 2.

Out of 194 MDR GNB isolates, 45% were isolated from patients admitted to the intensive care unit, 83.4% were isolated from patients previously exposed to different antimicrobial agents, 28% were isolated from patients under mechanical ventilation and/or urinary catheterization, and 34.7% were isolated from patients with hospital-acquired pneumonia and hospital-acquired infection Table 3.

**Table 3.**
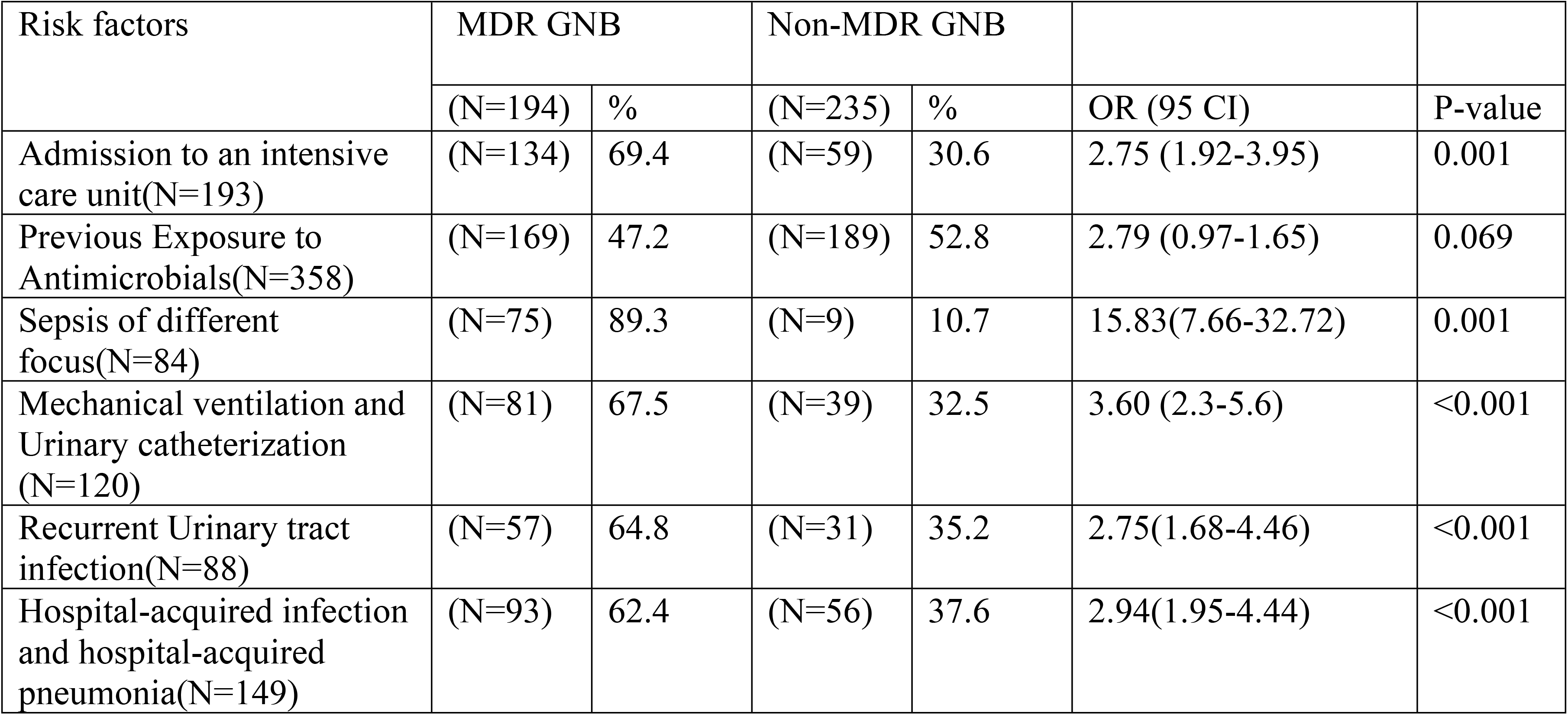
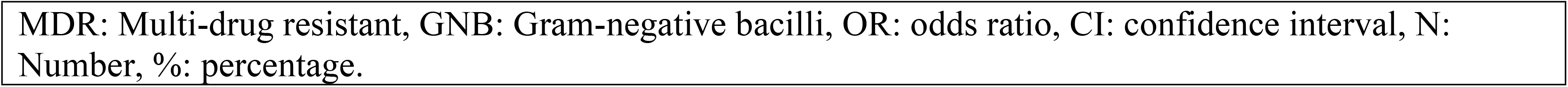
Univariate analysis of MDR GNB infection.

| Risk factors | MDR GNB |  | Non-MDR GNB |  | OR (95 CI) | P-value |
| --- | --- | --- | --- | --- | --- | --- |
|  | (N=194) | % | (N=235) | % |  |  |
| Admission to an intensive care unit(N=193) | (N=134) | 69.4 | (N=59) | 30.6 | 2.75 (1.92-3.95) | 0.001 |
| Previous Exposure to Antimicrobials(N=358) | (N=169) | 47.2 | (N=189) | 52.8 | 2.79 (0.97-1.65) | 0.069 |
| Sepsis of different focus(N=84) | (N=75) | 89.3 | (N=9) | 10.7 | 15.83(7.66-32.72) | 0.001 |
| Mechanical ventilation and Urinary catheterization (N=120) | (N=81) | 67.5 | (N=39) | 32.5 | 3.60 (2.3-5.6) | <0.001 |
| Recurrent Urinary tract infection(N=88) | (N=57) | 64.8 | (N=31) | 35.2 | 2.75(1.68-4.46) | <0.001 |
| Hospital-acquired infection and hospital-acquired pneumonia(N=149) | (N=93) | 62.4 | (N=56) | 37.6 | 2.94(1.95-4.44) | <0.001 |

MDR: Multi-drug resistant, GNB: Gram-negative bacilli, OR: odds ratio, CI: confidence interval, N: Number, %: percentage.

Admission to an intensive care unit (OR:2.75, 95% CI: 1.92-3.95, P value: 0.001), mechanical ventilation as source of infection and/or urinary catheterization as source of infection (OR:3.6, 95% CI :2.3-5.6, P value:<0.001), Hospital-acquired infection and/or hospital-acquired pneumonia as a source of infection (OR:2.94, 95% CI : 1.95-4.44, P value :<0.001), and recurrent urinary tract infection (OR:2.75, 95% CI:1.68-4.46, P value: <0.001), and sepsis originating from different focus (OR:15.83, 95% CI :7.66-32.72, P value:0.001) were significantly associated with acquiring multi-drug resistant gram-negative bacilli Table 3.

Out of 23 carbapenemase-positive organisms, 56.5% (13/23), 26.1% (6/23), 8.7% (2/23),8.7% (2/23) were isolated from urine, pus, blood, and tracheal aspirate respectively Table 4. Out of 23 carbapenemase-positive organisms,82.6% (19/23), 17.4% (4/23) were isolated from the patients admitted to the intensive care unit and unknown ward respectively Table 4. In this, study the prevalence of carbapenem-non susceptible and carbapenemase-producing gram-negative bacilli is 15.4% (66/429) and 5.4% (23/429) respectively Table 4.

**Table 4.** Distribution of carbapenemase among wards and specimen sources.

| Specimen types | Carbapenemase |  |  | Wards | Carbapenemase |  |  |
| --- | --- | --- | --- | --- | --- | --- | --- |
|  | Positive | Negative | Total |  | Positive | Negative | total |
| Blood | 2 | 7 | 9 | Emergency | 0 | 6 | 6 |
| Pleural fluid | 0 | 1 | 1 | ICU | 19 | 22 | 41 |
| Pus | 6 | 11 | 17 | Inpatient | 0 | 3 | 3 |
| CSF | 0 | 2 | 2 | outpatient | 0 | 2 | 2 |
| Tracheal aspirate | 2 | 3 | 5 | Unspecified | 4 | 10 | 14 |
| tissue | 0 | 1 | 1 | Total | 23 | 43 | 66 |
| urine | 13 | 18 | 31 |  |  |  |  |
| total | 23 | 43 | 66 |  |  |  |  |
| ICU-Intensive care unit, CSF-Cerebrospinal Fluid |  |  |  |  |  |  |  |

### Antimicrobial resistance profile of *Enterobacteriaceae*

Out of 429 gram-negative isolates, 293 were *Enterobacteriaceae* family (*E. coli, K. Pneumoniae, K. oxytoca, K. ozaenae)* Table 2. The highest resistance was observed against Ampicillin by *E. coli* [89.3%] Table 5. *K. Pneumoniae, K. oxytoca*, and *K. ozaenae* were intrinsically resistant to ampicillin hence not tested against them [12].

**Table 5.** Antibiotic susceptibility pattern of gram-negative bacilli against different antimicrobial classes.

|  |  | *E. coli* | *K. pneumoniae* | *K. oxytoca* | *K. ozaenae* | *E. cloacae* | *P. mirabilis* | *P. vulgaris* | *M.morganii* | *Acinetobacter species* | *P. aeruginosa* |
| --- | --- | --- | --- | --- | --- | --- | --- | --- | --- | --- | --- |
| Ampicillin & β-Lactam combinations | AMP %R | 89.3 | NA* | NA* | NA* | NA* | 100 | NA* | NA* | NA* | NA* |
|  | AMC %R | 33.8 | 64.4 | 33.3 | 87.5 | NA* | 38 | 0 | NA* | NA* | NA* |
|  | TPZ %R | 13.3 | 33.3 | 14.3 | 66.7 | 42.9 | 30 | 0 | 17 | 85.4 | 12.5 |
| 1st and 2nd generation cephalosporins | CZ %R | 60.4 | 85.7 | 57.1 | 100 | NA* | 80 | NA* | NA* | NA* | NA* |
|  | CXM %R | 64.1 | 91.4 | 60 | 100 | 66.7 | 71 | NA* | NA* | NA* | NA* |
| Extended-spectrum cephalosporins | CRO %R | 63.8 | 88.7 | 50 | 100 | 100 | 69 | 0 | 67 | 94.7 | NA* |
|  | CTX %R | 68.6 | 82.8 | 60 | 100 | 25 | 68 | 0 | 67 | 95.7 | NA* |
|  | CAZ %R | 65 | 81.1 | 33.3 | 100 | 71.4 | 83 | 0 | 0 | 93 | 14.3 |
|  | FEP %R | 55.7 | 81.8 | 40 | 85.7 | 75 | 89 | 0 | 0 | 86.1 | 21.7 |
| Carbapenems | IPM %R | 0 | 6.2 | 0 | 50 | 50 | 20 | 0 | 0 | 29.4 | 0 |
|  | MEM %R | 2.2 | 5.2 | 0 | 20 | 9.1 | 14 | 0 | 25 | 48.1 | 19.4 |
| Folate Pathway antagonists | SXT %R | 76.7 | 82.7 | 66.7 | 100 | 81.8 | 88 | 66.7 | 100 | 68.4 | NA* |
| Quinolone and Fluoroquinolone | NA %R | 83.3 | 80 | NA* | NA* | 0 | 100 | 0 | 100 | NA* | NA* |
|  | CIP %R | 62.9 | 62.3 | 25 | 87.5 | 66.7 | 71 | 50 | 60 | 77.6 | 21.9 |
| Tetracycline | TET %R | 84.8 | 60 | 66.7 | 60 | 50 | NA* | NA* | 0 | 92.9 | 0 |
| Nitrofurantoin | FM %R | 3.1 | 46.4 | 33.3 | 75 | 0 | NA* | NA* | NA* | NA* | NA* |
| Aminoglycosides | GEN %R | 20.5 | 71.2 | 0 | 66.7 | 60 | 25 | 0 | 33 | 72.7 | 0 |
|  | TOB %R | 33.6 | 37.2 | 25 | 72.7 | 57.1 | 63 | 0 | 25 | 54.3 | 17.2 |
|  | AMK %R | 3.1 | 11.1 | 0 | 12.5 | 0 | 46 | 0 | 0 | 29.5 | 4.3 |

Abbreviations: AMP-Ampicillin, AMC-Amoxicillin-clavulanate, TPZ-Piperacillin-tazobactam, CZ-Cefazolin, CXM-Cefuroxime, CRO-Ceftriaxone, CTX-Cefotaxime, CAZ-Ceftazidime, FEP-Cefepime, IPM-Imipenem, MEM-Meropenem, SXT-Trimethoprim-sulfamethoxazole, NA- Nalidixic acid, CIP-Ciprofloxacin, TET-Tetracycline, FM-Nitrofurantoin, GEN-Gentamycin, TOB-Tobramycin, AMK-Amikacin, %R-Percent resistant, NA\*-Not Applicable, 0-Zero resistance.

*K. ozaenae* showed 100% resistance to first, second, and third generation cephalosporins, as well as Trimethoprim-sulfamethoxazole, and *E. cloacae* showed 100% resistance to ceftriaxone.The overall resistance profile of *Enterobacteriaceae* to extended-spectrum cephalosporins ranges from Ceftriaxone [67.7%], Cefotaxime [73.6%], ceftazidime [73.6%], and Cefepime [66.5%]. *E. coli* showed high resistance to extended-spectrum cephalosporins; ceftriaxone [63.8%], cefotaxime [68.6%], Ceftazidime [65%] and cefepime [55.7%] Table 5. *K. Pneumoniae* showed high resistance to extended-spectrum cephalosporins; ceftriaxone [88.7%], cefotaxime [82.8%], ceftazidime [81.1%], cefepime [81.8%].

*E. coli* showed high resistance to ciprofloxacin [62.9%] and Trimethoprim-sulfamethoxazole [76.7%], likewise *K. Pneumoniae* showed high resistance to ciprofloxacin [62.3%] and Trimethoprim-sulfamethoxazole [82.7%] Table 5. The overall prevalence of *Enterobacteriaceae* resistance to gentamycin, tobramycin, amikacin is 41.9%, 37.5%, and 6.1% respectively. The overall prevalence of meropenem and imipenem resistance among the *Enterobacteriaceae* family was 4.2% and 6.5% respectively.

### Antimicrobial resistance profile of *Morganellacea* family

*M*.*morganii, P. mirabilis*, and *P. vulgaris* were reorganized to *Morganellacea* families [13]. A total of 26 *Morganellacea* families were isolated during the study period Table 2. *M*.*morganii* and *P. vulgaris* were intrinsically resistant to ampicillin, cefazolin, and cefuroxime and were not reported in Table 5. *M*.*morganii, P*.*mirabilis*, and *P*.*vulgaris* intrinsically to Antimicrobials were summarized in Table 5 as Not applicable [NA].

P. mirabilis showed 100% resistance to Ampicillin, Nalidixic acid, and Trimethoprim-sulfamethoxazole, likewise M. morganii showed 100% resistance to Nalidixic acid and Trimethoprim-sulfamethoxazole. family showed high resistance percentage to ceftriaxone [80.8%], cefotaxime [61%], ceftazidime [57.7%], and cefepime [63.6%]. The percentages of Meropenem and Imipenem resistance were 14% and 14.3% respectively Table 5. The Antimicrobial susceptibility patterns of the *Morganellacea* family were summarized in Table 5. The percentage of *Morganellacea* family resistance among aminoglycosides ranges from 26.3% to 47.5% Table 5.

### Antimicrobial resistance profile of *Acinetobacter* species

*Acinetobacter* species shows the high percentage of resistance to cefotaxime [95.7%], ceftriaxone [94.7], ceftazidime [93%], tetracycline [92.9%], and cefepime [86.1%]. *Acinetobacter* species showed lowest percentage of resistance to Amikacin [29.4%], Meropenem [48.1%], Imipenem [29.4%] Table 5.

#### Antimicrobial susceptibility patterns of *P. aeruginosa*

A total of 36 *P. aeruginosa* isolates were isolated during the study period. The highest percentage of resistance by *P. aeruginosa* was observed against Ciprofloxacin [N=32, 21.9%], Cefepime [N=32, 21.7%], and Meropenem [N=31, 19.4%] Table 5.

## Discussion

The prevalence of XDR, MDR, Carbapenemase-producing, and Carbapenem-resistant GNB is increasing [5, 6, 19]. In our study, the prevalence of MDR, XDR, Carbapenemase-producing, and carbapenem non-susceptible GNB is high. The most frequently isolated XDR organisms were *Acinetobacter* species 32.4% [24/74] which disagree with the study findings of Beyene et al (*E. coli* 18.1% was the dominant XDR GNB followed by *K. pneumoniae* 11.1% [19] and Gashaw et al (*Klebsiella* species 43.3% was the dominant XDR GNB) [20], the difference could be attributed to geographical differences, the number of samples studied, or the types of gram-negative bacteria considered. In the present study, the highest number of XDR organisms were recovered from urine samples 48.48% [16/33] and patients admitted to an intensive care unit 63.64% [21/33].

In the present study, the prevalence of XDR gram-negative bacilli was 7.7% [33/429], which is slightly lower than study at Ethiopian Public Health Institute, Ethiopia by Beyene et al 8.8% [19] the variation might be due to the fact the investigators analyzed only Enterobacterales, much lower than the findings from the study at Jimma, Ethiopia by Gashaw et al 41.3% [20], Arsho Advanced Medical Laboratory, Addis Ababa, Ethiopia by Bitew et al 34.3% [21], Tertiary Care Hospital, Pakistan by Abbas et al 64% [22]. The variation might be due to geographic location, the technique utilized, XDR definition, types of organism, etc.

In our study, the prevalence of Carbapenemase-producing gram-negative bacilli was 5.4%, which is higher than the prevalence of study conducted at the University of Gondar, Ethiopia by Eshetie et al 2.72% [23] and Ethiopian Public health Institute, Ethiopia by Beyene et al 2% [19]. However, our result was lower than the result of the study conducted at Tikur Anbessa specialized hospital, Ethiopia by Melese et al 12.12% [24], Three Hospitals in Amhara region, Ethiopia by Moges et al 15.7% [25], Felegehiwot Hospital, Ethiopia by Moges et al 16.2% [26], Sidama, Ethiopia by Alemayehu et al 9% [27], Mulago National Referral Hospital, Uganda by Okoche et al 22.4% [28], and data from laboratories in Spain by Lopez-Hernandez et al 62% [29]. The variation might be due to the technique utilized, modified carbapenem inactivation method was utilized in our investigation unlike other investigators who used the modified Hodge test, the number of bacterial isolates analyzed, geographic location. However, our study findings are in line with Prospective Cross-Sectional Study conducted at Felegehiwot Hospital, Ethiopia by Alebel et al 5.2% [30]. The dominant carbapenemase-producing gram-negative bacilli were *Klebsiella Pneumoniae* 8.3% [9/109] followed by *Acinetobacter* species 6.8% [5/74], the study results marginally coincides with the findings of the following researchers which demonstrated the dominant prevalence of carbapenemase-producing *Klebsiella pneumoniae*: Beyene et al *Klebsiella pneumoniae* 5.6% [19], Melese et al *Klebsiella pneumoniae* 10.5%[24], Moges et al *Klebsiella pneumoniae* 5.8%[25], Moges et al *Klebsiella pneumoniae* 10.1%[26], Lopez-Hernandez *Klebsiella pneumoniae* 45% [29]. The present study disagrees with finding by Okoche et al, which showed *E. coli* as the highest carbapenemase-producing organisms [28], The discrepancy could be explained by the fact that they utilized Boronic acid-based inhibition, a modified Hodge and EDTA double combination disk test, the number of samples analyzed, and a different geographical location., etc.

The highest number of carbapenemase producing isolates were recovered from urine samples 13/23(56.5%), which strongly disagrees with the results of study conducted by Moges et al in which the highest number of carbapenemase producing isolates was recovered from blood samples 22/24(91.6%)[25],

The prevalence of MDR was 45.2% which is much lower than the study conducted at Ethiopian Public Health Institute, Ethiopia by Beyene et al 94.5% [19], three referral hospitals, Ethiopia by Moges et al 85.8% [25], Felegehiwot Hospital, Ethiopia by Moges et al 80% and by Alebel et al 81.1% [26,30], Addis Ababa, Ethiopia by Teklu et al 68.3% [31], North Iran by Hemmati 62.8% [32], and Jimma medical center, Ethiopia by Biset et al 56.67% [33]. The variation might be due to the study population, the number of isolates assessed, the technique utilized, etc. However, our study findings are in line with the study conducted at Arsho Advanced Medical Laboratory, Addis Ababa, Ethiopia by Bitew et al 42.1% [21]. The dominant MDR isolates were *K. Pneumoniae* 80(73.4%) however our result disagrees with study findings of Beyene et al in which *E. coli* (99.3%)is the dominant one followed by *K*.*pneumoniae* (90.3%) [19] the observed variation could be attributable to the study period and proportion of bacterial isolates, Alebel et al in which *Acinetobacter species*(100%), *P. aeruginosa*(100%), *Citrobacter* species(100%), and *Enterobacter* cloacae(100%) were the dominant MDR isolates[30], the variation could be attributable to the fact that there were fewer samples evaluated, that samples were taken from intensive care unit patients, or that the study was conducted in a different geographic location and Biset et al [33] the difference might be since they only analyzed urine samples among pregnant women. Likewise, the following researchers reported the dominant prevalence of MDR Klebsiella *species* Moges et al *Klebsiella species (30*.*6%)* [25], Moges et al. *K. pneumoniae* (53.3%) [26] and, Teklu et al *K. pneumoniae* (83.5%) [31].

The prevalence of carbapenem carbapenem-resistant gram-negative bacilli is, 10.7% (46/429), in which *Acinetobacter* species account for about 39 percent of carbapenem-resistant isolates, the finding contradicts the study findings of the following researchers: Beyene et al 1.7% [19], Alebel et al 21% [30], Teklu et al 5.2% [31], Gashaw et al 25% [20], Melese et al 12.2% [24], in *K. pneumoniae* accounts for 100%, 40%, 50%,36%, and 75% of the gram-negative bacilli isolates respectively, the observed difference could be attributed to the types of gram-negative bacteria analyzed, as most of them only included *Enterobacteriaceae*, the techniques used, geographical location, and so on.

The total resistance profile of Enterobacterales to extended-spectrum cephalosporins ranges from 57.7% to 80.8%, which marginally agrees with results of study conducted by Beyene et al. 73.5% to 73.9% [19], but higher than the study findings of Teklu et al (60.3% to 62.2%) [31] and Moges et al (60.3% to 66%, in 2021) [25] lower than the study finding by Moges et al (87.2% to 96.6%, in 2019) [26], The geographical location, study time, and quantity of samples analyzed could all be factors in the reported discrepancy. Extended spectrum cephalosporin resistance was found in 17.5% of clinical Enterobacterales isolates studied in North America and Europe between 2016 and 2018 [40], the observed discrepancy could be attributed to the study’s large geographical scope, infection control techniques used in those settings, the number of samples tested, and so on.

## Conclusion and Recommendations

The finding from this study revealed a high prevalence of Multidrug-resistant (MDR), Extremely drug-resistant (XDR), carbapenemase-producing gram-negative bacteria, particularly among Intensive care unit (ICU) patients at the health facility level This necessitates a robust laboratory-based antimicrobial resistance monitoring and infection prevention and control program.

## Limitations of the study

The responsible genes for carbapenemase production were not genotypically assessed. Possible patient clinical impact was not assessed. Tigecycline, colistin, and Fosfomycin were not available and not used for AST.

## Acknowledgments

My deepest appreciation would go to Ethiopian Public Health institute for their support of reagents and supplies to conduct this research, and Dr. Adane Bitew and Dr. Eyasu Tigabu for their commitment in reviewing this manuscript.

## References

1. Ventola CL. The Antimicrobial resistance crisis: part 1: causes and threats. Pharmacy and therapeutics. 2015 Apr;40(4):277.

2. Aminov RI. A brief history of the antimicrobial era: lessons learned and challenges for the future. Frontiers in microbiology. 2010 Dec 8; 1:134.

3. Rolain JM, Canton R, Cornaglia G. Emergence of Antimicrobial resistance: the need for a new paradigm. Clinical Microbiology and Infection. 2012 Jul 1;18(7):615–6.

4. Watkins RR, Bonomo RA. Overview: global and local impact of antimicrobial resistance. Infectious Disease Clinics. 2016 Jun 1;30(2):313–22.

5. Souza GL, Rocha RF, Carvalho HD, Oliveira CD, Leite EM, Silva EU, Giarola LG, Couto BR, Starling CE. 2475. Incidence of Multidrug-Resistant, Extensively Drug-Resistant and Pandrug-Resistant Gram-Negative Bacteria in Brazilian Intensive Care Units. InOpen Forum Infectious Diseases 2019 Oct 2 (Vol. 6).

6. Kieffer N, Nordmann P, Aires-de-Sousa M, Poirel L. High prevalence of carbapenemase-producing Enterobacteriaceae among hospitalized children in Luanda, Angola. Antimicrobial agents and chemotherapy. 2016 Oct 1;60(10):6189–92.

7. Zhou L, Feng S, Sun G, Tang B, Zhu X, Song K, Zhang X, Lu H, Liu H, Sun Z, Zheng C. Extensively drug-resistant Gram-negative bacterial bloodstream infection in hematological disease. Infection and drug resistance. 2019; 12:481.

8. Magiorakos AP, Srinivasan A, Carey RT, Carmeli Y, Falagas MT, Giske CT, Harbarth S, Hindler JT, Kahlmeter G, Olsson-Liljequist B, Paterson DT. Multidrug-resistant, extensively drug-resistant and pan drug-resistant bacteria: an international expert proposal for interim standard definitions for acquired resistance. Clinical microbiology and infection. 2012 Mar 1;18(3):268–81.

9. Walther-Rasmussen J, Høiby N. OXA-type carbapenemases. Journal of Antimicrobial Chemotherapy. 2006 Mar 1;57(3):373–83.

10. Papp-Wallace KM, Bethel CR, Distler AM, Kasuboski C, Taracila M, Bonomo RA. Inhibitor resistance in the KPC-2 β-lactamase, a preeminent property of this class A β-lactamase. Antimicrobial agents and chemotherapy. 2010 Feb 1;54(2):890–7.

11. Raman G, Avendano EE, Chan J, Merchant S, Puzniak L. Risk factors for hospitalized patients with resistant or multidrug-resistant Pseudomonas aeruginosa infections: a systematic review and meta-analysis. Antimicrobial Resistance & Infection Control. 2018 Dec;7(1):1–4.

12. CLSI. Performance Standards for Antimicrobial Susceptibility Testing. 30th ed. CLSI supplement M100. Wayne, PA: Clinical and Laboratory Standards Institute; 2020.

13. CLSI. Performance Standards for Antimicrobial Susceptibility Testing. 22th ed. CLSI supplement M100. Wayne, PA: Clinical and Laboratory Standards Institute; 2012.

14. van der Zwaluw K, de Haan A, Pluister GN, Bootsma HJ, de Neeling AJ, Schouls LM. The carbapenem inactivation method (CIM), a simple and low-cost alternative for the Carba NP test to assess phenotypic carbapenemase activity in gram-negative rods. PloS one. 2015 Mar 23;10(3):e0123690.

15. Pierce VM, Simner PJ, Lonsway DR, Roe-Carpenter DE, Johnson JK, Brasso WB, Bobenchik AM, Lockett ZC, Charnot-Katsikas A, Ferraro MJ, Thomson RB. Modified carbapenem inactivation method for phenotypic detection of carbapenemase production among Enterobacteriaceae. Journal of clinical microbiology. 2017 Aug 1;55(8):2321–33.

16. CLSI. Performance Standards for Antimicrobial Susceptibility Testing. 27th ed. CLSI supplement M100. Wayne, PA: Clinical and Laboratory Standards Institute; 2017.

17. Jing X, Zhou H, Min X, Zhang X, Yang Q, Du S, Li Y, Yu F, Jia M, Zhan Y, Zeng Y. The simplified carbapenem inactivation method (sCIM) for simple and accurate detection of carbapenemase-producing Gram-negative bacilli. Frontiers in microbiology. 2018 Oct 30; 9:2391.

18. Use I, Of C, Packages THE, Samples K, Procedure T, Results ETHE. MIC Test Strip Technical Sheet KPC. 2015;7–8.

19. Beyene D, Bitew A, Fantew S, Mihret A, Evans M. Multidrug-resistant profile and prevalence of extended-spectrum β-lactamase and carbapenemase production in fermentative Gram-negative bacilli recovered from patients and specimens referred to National Reference Laboratory, Addis Ababa, Ethiopia. PloS one. 2019 Sep 25;14(9):e0222911.

20. Gashaw M, Berhane M, Bekele S, Kibru G, Teshager L, Yilma Y, Ahmed Y, Fentahun N, Assefa H, Wieser A, Gudina EK. Emergence of high drug resistant bacterial isolates from patients with health care associated infections at Jimma University medical center: a cross sectional study. Antimicrobial Resistance & Infection Control. 2018 Dec;7(1):1–8.

21. Bitew A, Tsige E. High Prevalence of Multidrug-Resistant and Extended-Spectrum β-Lactamase-Producing Enterobacteriaceae: A Cross-Sectional Study at Arsho Advanced Medical Laboratory, Addis Ababa, Ethiopia. Journal of tropical medicine. 2020 Apr 30;2020.

22. Abbas S, Sabir AU, Khalid N, Sabir S, Khalid S, Haseeb S, Khan MN, Ajmal WM, Azhar F, Saeed MT. Frequency of Extensively Drug-Resistant Gram-Negative Pathogens in a Tertiary Care Hospital in Pakistan. Cureus. 2020 Dec;12(12).

23. Eshetie S, Unakal C, Gelaw A, Ayelign B, Endris M, Moges F. Multidrug resistant and carbapenemase producing Enterobacteriaceae among patients with urinary tract infection at referral Hospital, Northwest Ethiopia. Antimicrobial resistance and infection control. 2015 Dec;4(1):1–8.

24. Legese MH, Weldearegay GM, Asrat D. Extended-spectrum beta-lactamase-and carbapenemase-producing Enterobacteriaceae among Ethiopian children. Infection and drug resistance. 2017; 10:27.

25. Moges F, Gizachew M, Dagnew M, Amare A, Sharew B, Eshetie S, Abebe W, Million Y, Feleke T, Tiruneh M. Multidrug resistance and extended-spectrum beta-lactamase producing Gram-negative bacteria from three Referral Hospitals of Amhara region, Ethiopia. Annals of clinical microbiology and antimicrobials. 2021 Dec;20(1):1–2.

26. Moges F, Eshetie S, Abebe W, Mekonnen F, Dagnew M, Endale A, Amare A, Feleke T, Gizachew M, Tiruneh M. High prevalence of extended-spectrum beta-lactamase-producing gram-negative pathogens from patients attending Felege Hiwot Comprehensive Specialized Hospital, Bahir Dar, Amhara region. PloS one. 2019 Apr 15;14(4):e0215177.

27. Alemayehu T, Asnake S, Tadesse B, Azerefegn E, Mitiku E, Agegnehu A, Nigussie N. Phenotypic Detection of Carbapenem-Resistant Gram-Negative Bacilli from a Clinical Specimen in Sidama, Ethiopia: A Cross-Sectional Study. Infection and Drug Resistance. 2021;14:369.

28. Okoche D, Asiimwe BB, Katabazi FA, Kato L, Najjuka CF. Prevalence and characterization of carbapenem-resistant Enterobacteriaceae isolated from Mulago National Referral Hospital, Uganda. PLoS One. 2015 Aug 18;10(8):e0135745.

29. López-Hernández I, Delgado-Valverde M, Fernández-Cuenca F, López-Cerero L, Machuca J, Pascual Á. Carbapenemase-Producing Gram-Negative Bacteria in Andalusia, Spain, 2014–2018. Emerging infectious diseases. 2020 Sep;26(9):2218.

30. Alebel M, Mekonnen F, Mulu W. Extended-Spectrum β-Lactamase and Carbapenemase Producing Gram-Negative Bacilli Infections Among Patients in Intensive Care Units of Felegehiwot Referral Hospital: A Prospective Cross-Sectional Study. Infection and Drug Resistance. 2021; 14:391.

31. Teklu DS, Negeri AA, Legese MH, Bedada TL, Woldemariam HK, Tullu KD. Extended-spectrum beta-lactamase production and multi-drug resistance among Enterobacteriaceae isolated in Addis Ababa, Ethiopia. Antimicrobial Resistance & Infection Control. 2019 Dec;8(1):1–2.

32. Hemmati H, Hasannejad-Bibalan M, Khoshdoz S, Khoshdoz P, Kalurazi TY, Ebrahim-Saraie HS, Nalban S. Two years study of prevalence and antibiotic resistance pattern of Gram-negative bacteria isolated from surgical site infections in the North of Iran. BMC research notes. 2020 Dec;13(1):1–6.

33. Biset S, Moges F, Endalamaw D, Eshetie S. Multi-drug resistant and extended-spectrum β-lactamases producing bacterial uropathogens among pregnant women in Northwest Ethiopia. Annals of Clinical Microbiology and Antimicrobials. 2020 Dec; 19:1–9.

34. Gashe F, Mulisa E, Mekonnen M, Zeleke G. Antimicrobial Resistance Profile of Different Clinical Isolates against Third-Generation Cephalosporins. Journal of pharmaceutics. 2018 Sep 9; 2018:5070742-.

35. Breurec S, Bouchiat C, Sire JM, Moquet O, Bercion R, Cisse MF, Glaser P, Ndiaye O, Ka S, Salord H, Seck A. High third-generation cephalosporin resistant Enterobacteriaceae prevalence rate among neonatal infections in Dakar, Senegal. BMC infectious diseases. 2016 Dec;16(1):1–7.

36. Patolia S, Abate G, Patel N, Patolia S, Frey S. Risk factors and outcomes for multidrug-resistant Gram-negative bacilli bacteremia. Therapeutic advances in infectious disease. 2018 Jan;5(1):11–8.

37. Nseir S, Di Pompeo C, Cavestri B, Jozefowicz E, Nyunga M, Soubrier S, Roussel-Delvallez M, Saulnier F, Mathieu D, Durocher A. Multiple-drug–resistant bacteria in patients with severe acute exacerbation of chronic obstructive pulmonary disease: Prevalence, risk factors, and outcome. Critical care medicine. 2006 Dec 1;34(12):2959–66.

38. Huang SF, Chang JS, Sheu CC, Liu YT, Lin YC. An antibiotic decision-making tool for patients with pneumonia admitted to a medical intensive care unit. International journal of antimicrobial agents. 2016 Sep 1;48(3):286–91.

39. Chen G, Xu K, Sun F, Sun Y, Kong Z, Fang B. Risk Factors of Multidrug-Resistant Bacteria in Lower Respiratory Tract Infections: A Systematic Review and Meta-Analysis. Canadian Journal of Infectious Diseases and Medical Microbiology. 2020 Jun 30;2020.

40. Belley A, Morrissey I, Hawser S, Kothari N, Knechtle P. Third-generation cephalosporin resistance in clinical isolates of Enterobacterales collected between 2016–2018 from USA and Europe: genotypic analysis of β-lactamases and comparative in vitro activity of cefepime/enmetazobactam. Journal of global antimicrobial resistance. 2021 Jun 1;25:93–101.

